# Characterization of the Preclinical in Vitro and in Vivo Pharmacokinetic Properties of the Cyclic Oligomeric Depsipeptide Ryanodine Receptor 2 Inhibitor *ent*-Verticilide

**DOI:** 10.64898/2026.10.26.761690

**Authors:** Tri Q. Do, Rachel D. Crouch, Kevin Wu, Madelaine P. Thorpe, Abigail N. Smith, Jeffrey N. Johnston, Wendell S. Akers, Björn C. Knollmann

**Affiliations:** Vanderbilt Center for Arrhythmia Research and Therapeutics, Division of Clinical Pharmacology and Genetic Medicine, Department of Medicine, Vanderbilt University Medical Center, Nashville, TN, United States; Center for Drug Metabolism and Pharmacokinetics, Pharmaceutical Sciences Research Center, Lipscomb University College of Pharmacy, Nashville, TN, United States; Department of Chemistry, Vanderbilt Institute of Chemical Biology, Vanderbilt University, Nashville, TN, United States; Department of Pharmacology, Vanderbilt University School of Medicine, Nashville, TN, United States

**Keywords:** Cardiac arrhythmia, Cyclic oligomeric depsipeptides Ryanodine receptor 2, Drug metabolism and pharmacokinetics

## Abstract

*ent*-Verticilide is a synthetic cyclic oligomeric depsipeptide that inhibits ryanodine receptor 2 (RyR2)-mediated calcium (Ca) leak and has demonstrated antiarrhythmic activity in mice. This study characterized the preclinical absorption, distribution, metabolism, and pharmacokinetic properties of *ent*-verticilide and investigated factors affecting its oral exposure. *ent*-Verticilide was stable in rat plasma and simulated gastric and intestinal fluids but exhibited extensive plasma protein binding (>94%) and low apparent permeability in Caco-2 cells, with no evidence of pronounced net efflux. In rat and human liver microsomes, *ent*-verticilide underwent rapid NADPH-dependent metabolism, with an in vitro half-life of 5.43 min and 4.35 min, respectively, and a predicted hepatic extraction ratio of 0.90 and 0.92, respectively. Following intravenous administration in rats, *ent*-verticilide exhibited a systemic clearance of 25.5 mL/min/kg and a large steady-state volume of distribution (4.72 L/kg). Absolute oral bioavailability was approximately 16%. Simultaneous portal and systemic sampling estimated intestinal availability (*F_a_F_g_*) at 0.25 and hepatic availability (*F_h_*) at 0.64, suggesting that processes occurring before hepatic entry are a major limitation to oral exposure. Pretreatment with the CYP inhibitor 1-aminobenzotriazole increased portal and systemic exposure approximately 10.6- and 18.5-fold, respectively, and increased apparent *F_h_* to approximately 1. Collectively, these findings identify limited intestinal availability and CYP-mediated presystemic metabolism as important determinants of *ent*-verticilide exposure and provide a framework for its continued development as an RyR2-targeted therapeutic.

**SIGNIFICANT STATEMENT:** This study defines key factors governing the pharmacokinetic disposition of *ent*-verticilide, a RyR2 inhibitor with antiarrhythmic activity. *ent*-Verticilide exhibited low intestinal permeability, extensive protein binding, and rapid CYP-dependent metabolism. Portal and systemic pharmacokinetic studies identified limited intestinal availability as a major determinant of oral exposure, while CYP inhibition markedly increased drug exposure. These findings provide a mechanistic framework for understanding *ent*-verticilide disposition and inform strategies to improve its oral pharmacokinetic properties and further therapeutic development.

## INTRODUCTION

Ryanodine receptor 2 (RyR2) is the major intracellular calcium (Ca)-release channel in cardiac muscle and plays a central role in excitation–contraction coupling.^1^ Hyperactive RyR2 can result in Ca leak, promoting delayed afterdepolarizations and triggered electrical activity that contribute to cardiac arrhythmias.^2^ Consequently, pharmacological stabilization or inhibition of pathological RyR2-mediated Ca release has emerged as a potential strategy for cardiac arrhythmias.^2^

*ent*-Verticilide is the synthetic enantiomer of the cyclic oligomeric depsipeptide verticilide and represents a structurally distinct RyR2 inhibitor.^3,4^ Previous work demonstrated that *ent*-verticilide and its 18-membered ring analog *ent*-B1 inhibit RyR2-mediated Ca leak and suppress cardiac arrhythmia in mice,^3,5^ establishing cyclic oligomeric depsipeptides as a promising scaffold for the development of RyR2-targeted therapeutics. Despite its pharmacological activity, considerably less is known about the drug disposition properties of *ent*-verticilide. In particular, the physicochemical and structural features of a relatively large cyclic depsipeptide can present challenges for oral drug delivery, including extensive protein binding, limited intestinal permeability, nonspecific binding, and presystemic elimination.^6,7^ Defining these properties is therefore important for understanding the relationship between administered dose and pharmacologically relevant systemic exposure, and advancing *ent*-verticilide towards therapeutic development.

Accordingly, the present study aimed to characterize the absorption, distribution, and metabolic properties that govern the pharmacokinetic disposition of *ent*-verticilide. Its plasma protein binding, stability in biorelevant gastrointestinal media, intestinal permeability, and hepatic metabolic stability were evaluated using complementary in vitro approaches. In vivo pharmacokinetic studies were then conducted in Sprague–Dawley rats following intravenous and oral administration, with simultaneous hepatic portal vein (HPV) and jugular vein (JVC) sampling to assess presystemic and systemic exposure. Finally, the effect of cytochrome P450 (CYP) inhibition by 1-aminobenzotriazole (ABT) was examined to determine the contribution of CYP-mediated metabolism to the oral disposition of *ent*-verticilide. Together, these studies were intended to identify the major determinants of *ent*-verticilide exposure and provide a pharmacokinetic framework for its continued development as an RyR2-targeted antiarrhythmic agent.

## METHODS

### Chemicals and Reagents

Dimethyl sulfoxide (DMSO), magnesium chloride (MgCl_2_), monobasic potassium phosphate, dibasic potassium phosphate, carbamazepine, nicotinamide adenine dinucleotide phosphate tetrasodium salt (NADPH), atenolol, metoprolol, digoxin, verapamil hydrochloride, and warfarin were purchased from Sigma Aldrich (St. Louis, MO). The synthesis of *ent*-verticilide has been described elsewhere.^3^ Optima high performance liquid chromatography (HPLC)-grade acetonitrile and formic acid were purchased from Fisher Scientific. HPLC water was obtained from a Milli-Q plus purification system (Millipore, Bedford, MA). Drug-free rat and human plasma were purchased from BioIVT (Westbury, NY). TRANSIL PPB and HSA binding kits were purchased from Sovicell (Leipzig, Germany).

### Liver Fractions, Caco-2 Cells and Animals

Commercially available liver microsomes from rat (catalog number R1000 / lot number 2110178 / n = 428) and human (catalog number H2610 / lot number 1910096 / n = 200) were purchased from Sekisui XenoTech (Kansas City, KS). Rat liver microsomes were pooled from male donors and human liver microsomes were pooled from a mixture of male and female donors. Caco-2 cells were purchased from ReadyCell (Barcelona, Spain). In vivo pharmacokinetic studies were conducted by a contract research organization (Pharmaron, Inc.) in AAALAC-accredited animal facilities and followed IACUC-approved study protocols (Protocols: #PH-DMPK-VUMC-23-004, #PH-DMPK-VUMC-23-006).

### Plasma Protein Binding

Preliminary studies to evaluate plasma protein binding of *ent*-verticilide by rapid equilibrium dialysis were unsuccessful due to high non-specific binding of the compound to the polytetrafluoroethylene and polypropylene reservoir plates such that no *ent*-verticilide could be detected in the dialysate. A modified ultrafiltration technique used for compounds with high non-specific binding was able to detect a very low level of free *ent*-verticilide in plasma filtrate with an estimated plasma protein binding exceeding 99% (unpublished data). TRANSIL PPB and HSA assay kits (Sovicell, Inc.) utilizing purified plasma proteins immobilized on silica beads and ultralow binding tubes to minimize high nonspecific binding were used to further characterize the plasma protein binding properties of *ent*-verticilide.

The TRANSIL PPB binding kit evaluated the affinity of *ent*-verticilide to human serum albumin (HSA) and alpha1-acid glycoprotein (AAG) immobilized on silica beads at a physiologic ratio of 24:1. The TRANSIL HSA Binding kit evaluated the binding affinity of *ent*-verticilide to varying concentrations of HSA (3.1 – 58.8 mM) immobilized on silica beads. Plasma protein binding for each assay was determined by incubating a fixed concentration of *ent*-verticilide with varying concentrations of the plasma proteins immobilized on silica beads suspended in 10 mM phosphate buffered saline (PBS) adjusted to pH 7.4. Reference wells without plasma protein covered silica beads served as reference controls to account for non-specific binding in each assay. *ent*-Verticilide was tested in both assays at four different fixed concentrations (0.01 µM, 0.1 µM, 1 µM, and 10 µM) to evaluate concentration-dependent protein binding to albumin. A single concentration of warfarin (1 µM) was included as a positive control for high plasma protein binding to HSA. Compounds were prepared as 16x stock solutions in 10 mM PBS adjusted to pH 7.4 and 15 µL of each compound stock solution was added to a column of 8 wells for a total assay volume of 240 µL per well. The plate was incubated at room temperature for 12 minutes on a plate shaker at 1000 rpm then centrifuged for 10 minutes at 750 x g. The supernatant (50 µL) from each well was transferred into a standard 96-well plate containing acetonitrile with internal standard and analyzed by liquid chromatography tandem mass spectrometry (LC-MS/MS). The peak area response for supernatant samples from the reference and test wells represented the total and free concentrations for each compound in the assay, respectively. The dissociation constant (K_D_) for each compound was calculated using equation 1:

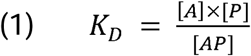

where [AP] represents concentration of compound bound to plasma protein determined by subtracting the free compound concentration from the total compound concentration, [A] is the free concentration of compound, and [P] denotes the protein concentration in each test well. The free concentration of compound can also be expressed using equation 2:

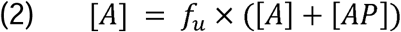

where *f*_u_ is the fraction unbound calculated by dividing the free concentration [A] by the total concentration ([A] + [AP]). When substituting equation 2 into equation 1 and rearranging, a linear equation was derived and used to model the data from the TRANSIL PPB and HSA Binding Kit with equation 3 to determine the dissociation constant by taking the inverse of the slope (1/K_D_):

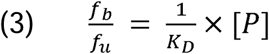

where *f*_b_ is the fraction bound calculated by subtracting *f*_u_ from 1. Estimation of the fraction bound to purified plasma proteins can be calculated using the total concentration of HSA and AAG in each assay along with the estimated K_D_ of the compound for each protein using equation 4:

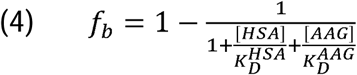

### Biorelevant Media Stability

The in vitro stability of *ent*-verticilide was evaluated in drug-free rat plasma and simulated gastrointestinal fluids. Fasted state simulated gastric fluid (FaSSGF; pH 1.6, in mM: chloride 59, sodium 34, taurocholate 0.08, phospholipids 0.02), and fasted state simulated intestinal fluid (FaSSIF; pH 6.5, in mM: chloride 106, sodium 148, phosphate 29, taurocholate 3, phospholipids 0.75) were prepared using Biorelevant 3F Powder®, corresponding Biorelevant Buffer Concentrates, and purified water according to manufacturer recommendations. A modified FaSSIF pH 1.6 was also prepared by adding concentrated hydrochloric acid to FaSSIF pH 6.5 to explore the impact of pH versus the presence of taurocholate and phospholipids on *ent*-verticilide stability relative to FaSSGF. Triplicate samples for each biorelevant media were preincubated in a low binding polypropylene 96-well plate for 30 minutes on a microplate shaker at 37 °C prior to the addition of *ent*-verticilide (final concentration 2500 ng/mL). Following the addition of *ent*-verticilide to each well, samples were mixed thoroughly with a pipette, and a 50 µL sample was removed at 0, 15, 30, 45, 60, 90, and 120 minutes to evaluate compound stability in each condition over time relative to the baseline sample at time 0. After the 120-minute sample was collected, an equal volume of fresh FaSSIF pH 6.0 containing taurocholate and phospholipids as surfactants was added to each well to evaluate the potential of nonspecific binding of *ent*-verticilide over time in each biorelevant media condition. Wells were incubated in the refrigerator for an additional 4 hours and a 50 µL aliquot was collected to evaluate the *ent*-verticilide concentration and recovery in each condition. All samples from the time course study and subsequent recovery experiment were immediately transferred to a clean 96-well plate containing ice-cold acetonitrile with an internal standard prior to LC-MS/MS analysis.

### Caco-2 Permeability Studies

Polarized human colorectal adenocarcinoma-derived (Caco-2) cells were differentiated (passage 55 to 65) for 21 days in 24 Transwell® inserts with a semiporous polystyrene membrane (0.4 µm; CORNING Cat#3378) at a density of 150,000 – 180,000 cells per well in Dulbecco’s Modified Eagle’s Medium (glucose 1 g/L, fetal bovine serum 10% v/v, L-glutamine 2 mM, penicillin 110 U/mL, and streptomycin 100 µg/mL). Transepithelial electrical resistance (TEER) measurements greater than 1000 Ω x cm^2^ between the apical and basal compartments were used to identify inserts with cell monolayers suitable for transport studies. Hank’s balanced salt solution (HBSS; pH 7.4) was used as the standard transport buffer for all permeability experiments. Final concentrations of test and reference compounds were prepared in HBSS and run in duplicate to assess permeability. Test compounds were diluted to a final concentration of 10 µM with less than 1% of DMSO in the assay buffer. Reference compounds (10 µM) used as positive controls included atenolol and metoprolol as low and high permeability compounds along with digoxin as a known substrate for the efflux transporter, p-glycoprotein.

Prior to starting transport experiments, cell culture media was gently aspirated from both sides of the cell monolayer inserts and washed twice by adding 250 µL and 750 µL of HBSS pre-warmed to 37 °C to the apical and basal compartment, respectively. Following the second wash, the plate was returned to the cell incubator (37 °C, 5% CO_2_) for 30 minutes. For apical-to-basal (A-B) permeability studies, test compounds prepared in pre-warmed HBSS (275 µL) were applied to the apical compartment (donor side) and each insert was immediately placed into the basal compartment (receiver side) containing pre-warmed HBSS (750 µL). For basal-to-apical (B-A) permeability studies, pre-warmed HBSS (250 µL) was applied to the apical compartment (receiver side) and each insert was immediately placed into the basal compartment containing test compounds (donor side) containing test compound in pre-warmed HBSS (775 µL).

A 25 µL sample was immediately removed from the donor side of the plate (apical for A-B studies; basal for B-A studies) at time zero (T_0_) and transferred to a separate 96-well plate containing 75 µL of acetonitrile with internal standard for LC-MS/MS analysis. Plates were returned to the cell incubator for 2 hours to allow mass transport of test and reference compounds to cross the cell monolayer from the donor side to the receiver side. A 25 µL sample was collected from both the donor and receiver side after 2 hours (T_2_) and transferred to a 96-well plate containing 75 µL of acetonitrile with internal standard for LC-MS/MS analysis. All samples collected during the transport assay (T_0_ and T_2_) were analyzed by a quantitative LC-MS/MS method to determine the initial donor concentration, overall compound recovery, and the bidirectional permeability (A-B and B-A) of each test and reference compound.

The permeability for each compound in Caco cells was calculated using the following equation:

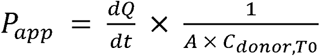

where the apparent permeability (*P_app_*; cm/sec) corresponds to the amount of compound that crosses the cell monolayer as function of time (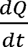; nmol/sec) divided by the product of the cell monolayer surface area (cm^2^) and the initial concentration of the compound in the donor compartment at time zero (*C_donor,To_*; nmol/mL).

An efflux ratio (ER) for the bidirectional transport of each compound across the cell monolayer was used as a general measure to evaluate the involvement of passive and active transport processes using the following equation:

The ER for each compound was calculated by dividing the *P_app_* in the B-A direction by the *P_app_*in the A-B direction:

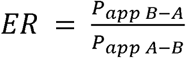

Where *P_app B—A_* and *P_app A—B_* refers to the permeability of each compound in the B-A and A-B directions, respectively.

Compound recovery for each transwell was determined using the following mass balance equation:

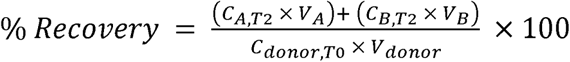

Where *C_A,T2_* and *C_B,T2_*correspond to compound concentrations at timepoint 2 hours in the apical and basal compartments, respectively. The term *C_donor,To_* is the initial concentration of the compound in the donor compartment at time zero. The terms *Y_A_*, *Y_B_*, and *Y_donor_*correspond to the volume of buffer in the apical, basal, and donor compartments, respectively.

Cell monolayer integrity for each well was evaluated at the end of experiments by measuring the A-B permeability of lucifer yellow. HBSS was gently aspirated from the basal compartment and replaced with fresh, pre-warmed HBSS (750 µL). Pre-warmed HBSS (250 µL) containing lucifer yellow (100 µM) was applied to the apical compartment (donor side) and each insert was immediately placed into the basal compartment (receiver side). Plates were protected from light and placed in the cell incubator (at 37 °C and 5% CO_2_) for 1 hour. At the end of the incubation period, a 200 µL sample was collected from the basal compartment and transferred to 96-well plate for fluorescence-based assays. Fluorescence intensity of samples and a standard curve prepared from the lucifer yellow working solution were measured in a fluorometer at 485/527 excitation/emission wavelengths. Lucifer yellow paracellular flux values ≤ 0.5 % were used as a strong indicator of cell barrier integrity.

### Metabolic Stability in Rat and Human Hepatic Microsomal Fractions

Intrinsic clearance (*Ci_int_*) of *ent*-verticilide was estimated in rat liver microsomes using the substrate depletion method. *ent*-Verticilide (1 µM) was incubated at 37 °C (shaking water bath) in a potassium phosphate buffered (100 mM, pH 7.4) mixture containing 3 mM MgCl_2_ and rat and human liver microsomes (0.5 mg/mL) in the presence or absence of NADPH (2 mM). Corresponding control incubations were conducted without NADPH. *ent*-Verticilide was evaluated at a final concentration of 1 µM. Verapamil was evaluated separately at 1 µM under otherwise identical conditions as a positive-control CYP3A substrate to verify NADPH-dependent metabolic activity. The final organic-solvent concentration was held constant within each experiment and did not exceed 0.6%. At 0, 3, 7, 15, 30, and 45 minutes, 100-µL aliquots were removed from each incubation and transferred immediately to a 96-well protein-precipitation filtration plate containing 300 µL of ice-cold acetonitrile supplemented with 50 nM carbamazepine as the internal standard. Precipitated proteins were removed by filtration into a polypropylene 96-well collection plate using a positive-pressure manifold. Filtered extracts were evaporated to dryness under a stream of nitrogen. The resulting residues were reconstituted in 100 µL of acetonitrile/water (30:70, v/v) before LC–MS/MS analysis.

Samples were introduced using a CTC Analytics PAL-xt DW autosampler, and 10 µL was injected into a Shimadzu LC-20AD series HPLC system coupled to a SCIEX 6500 QTRAP mass spectrometer (SCIEX, Foster City, CA, USA). The mass spectrometer was equipped with an electrospray ionization source (500 °C ion source temperature) operated in positive-ion mode (4500 V ion spray voltage). Chromatographic separation was performed using a Kinetex® C18 column (50 × 2.1 mm, 1.7-µm particle size; Phenomenex, Torrance, CA, USA) maintained at 60 °C. Mobile phase A consisted of 0.1% formic acid in ultrapure water, and mobile phase B consisted of 0.1% formic acid in acetonitrile. The mobile phases were delivered at a flow rate of 0.5 mL/min. The initial mobile-phase composition was 5% B. Mobile phase B was linearly increased from 5% to 90% over 0.5 minutes. The 90% B condition was maintained from 0.5 to 2.00 minutes, after which the composition was returned to 5% B from 2.00 to 2.01 minutes. The column was re-equilibrated at 5% B until the end of the 2.5-minute analytical run. *ent*-Verticilide was detected by multiple-reaction monitoring using the precursor-to-product-ion transition *m/z* 853.6→214.1. Carbamazepine (internal standard) was monitored using the transition *m/z* 237.1→194.0. Verapamil was monitored using the transition *m/z* 455.0→165.0. Peak areas were integrated using Analyst 1.7.0. The ion-source parameters, collision energies, and MRM dwell times for *ent*-verticilide, carbamazepine, and verapamil are reported in **Table 1**.

**Table 1:** In Vitro Plasma Protein Binding of *ent*-verticilide. Warfarin was included for reference.

| Compound | Concentration<br>($\mu\text{M}$) | Plasma Protein Binding | | Human Serum Albumin | |
| --- | --- | --- | --- | --- | --- |
|  |  | % Bound | % Free | % Bound | % Free |
| Warfarin | 1 | 97.1 | 2.9 | 98.4 | 1.6 |
| *ent*-Verticilide | 0.01 | 99.1 | 0.9 | 96.4 | 3.6 |
|  | 0.1 | 99.5 | 0.5 | 96.9 | 3.1 |
|  | 1 | 98.4 | 1.6 | 96.5 | 4.1 |
|  | 10 | 94.2 | 5.8 | 92.7 | 7.3 |

For each sample, the LC–MS/MS area ratio was calculated by dividing the analyte peak area by the carbamazepine peak area:

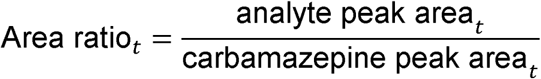

The percentage of *ent*-verticilide or verapamil remaining at each time point was calculated by normalizing the area ratio at that time point to the corresponding area ratio at 0 minutes:

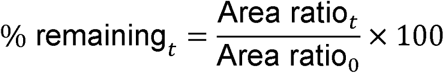

The percentage of parent compound remaining was natural-log transformed and plotted against incubation time. The first-order depletion rate constant, k, was determined as the negative of the slope of the log-linear regression:

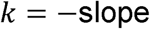

The in vitro half-life was calculated as:

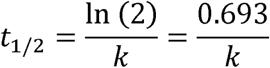

Measurements below 5% remaining substrate were excluded from regression analysis.

In vitro hepatic intrinsic clearance (*CL_int,opp,scołed_*, ml/min/kg) for each species was estimated from liver microsomes using the substrate depletion method according to the equation below:^8^

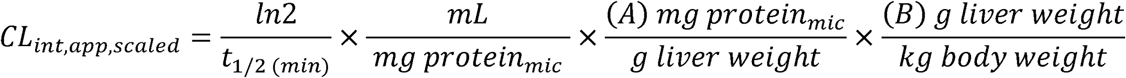

A microsomal value (mg protein per g liver weight) of A = 61 (rat) or 32 (human) was used for calculations, as previously reported.^9,10^ Values for B (g liver weight per kg body weight) were also derived from the literature; B = 40 and 25 for rat and human, respectively.^11^

Because binding within the liver microsomal incubations was not incorporated into the calculations, the resulting values were designated scaled apparent intrinsic clearance. Binding-unadjusted hepatic clearance was predicted using the well-stirred model:^12^

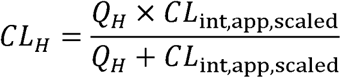

The corresponding hepatic extraction ratio was calculated as:

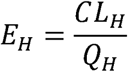

Species-specific hepatic blood-flow values of 70 and 21 mL/min/kg were used for rat and human, respectively.^11,13^

These calculations did not incorporate binding in plasma, blood, or the liver microsomal incubations. Accordingly, they were considered exploratory, binding-unadjusted predictions rather than definitive estimates of in vivo hepatic clearance. In the absence of binding terms, the model is mathematically equivalent to assuming that the entire measured drug concentration was available for hepatic metabolism. Although a blood-to-plasma concentration ratio of 0.85 was reported previously for *ent*-verticilide,^14^ that value was not incorporated because blood and plasma binding corrections were outside the scope of the current exploratory analysis.

### In Vivo Pharmacokinetic Studies

The in vivo pharmacokinetic properties of *ent*-verticilide were evaluated in rats following intravenous (IV) or oral (PO) dose administration. Preclinical formulations consisted of 10% Tween 20, 10% DMSO, 40% H_2_O, and 40% PEG400 with a final *ent*-verticilide concentration of 0.2 mg/mL (IV) or 0.9 mg/mL (PO). Pharmacokinetic parameters were evaluated in male Sprague Dawley rats administered either a single IV (1 mg/kg) or PO (9 mg/kg) dose of *ent*-verticilide (n=3 per treatment group). Individual rats were double cannulated to allow simultaneous blood sampling from both the portal (pre-systemic exposure) and jugular (systemic exposure) vein. Blood samples were serially collected at 0.083, 0.25, 0.5, 1, 3, 4, 8 and 24 hours following IV administration and at 0.25, 0.5, 1, 2, 4, 8, and 24 hours following PO administration. The impact of pre-systemic intestinal and hepatic metabolism on the oral bioavailability of *ent*-verticilide was evaluated in a separate pharmacokinetic study by pretreating rats (n=3 per treatment group) with the pan CYP450 inhibitor, ABT.^15^ Vehicle or ABT (100 mg/kg) was administered PO to rats 15 hours prior to a single PO dose of *ent*-verticilide (9 mg/kg). Blood samples were serially collected from both the portal vein and jugular vein at 0.25, 0.5, 1, 2, 4, 8, and 24 hours following *ent*-verticilide administration. Blood samples collected during the pharmacokinetic studies were centrifuged immediately to obtain plasma and stored at −80 °C until analysis by liquid chromatography coupled with tandem mass spectrometry (LC–MS/MS).

Individual plasma concentration–time profiles were analyzed by noncompartmental analysis using Phoenix WinNonlin version 8.3 (Certara USA, Inc., Princeton, NJ, USA). Nominal sample-collection times were used for all noncompartmental analysis. The maximum observed plasma concentration (*C_max_*) and time to reach the maximum observed concentration (*T_max_*) were obtained directly from the observed concentration–time data. The terminal elimination rate constant (*y_z_*) was estimated by log-linear regression of the terminal portion of each concentration–time profile. The terminal elimination half-life (*t_1/2_*) was calculated as:

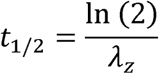

The area under the plasma concentration–time curve from time zero to the last quantifiable concentration (*AUC_O—t_*) was calculated using the linear-up/log-down trapezoidal method. The AUC from time zero extrapolated to infinity was calculated as:

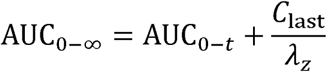

where *C_last_* is the last quantifiable concentration. The percentage of *AUC_O—e_* obtained by extrapolation was calculated for each profile to assess the reliability of the extrapolated AUC.

Following intravenous (IV) administration, the concentration extrapolated to time zero (*C_O_*) was estimated from the initial concentration–time data, as appropriate. Systemic clearance (*Ci*), terminal-phase volume of distribution (*Y_z_*), and steady-state volume of distribution (*Y_ss_*) were calculated as:

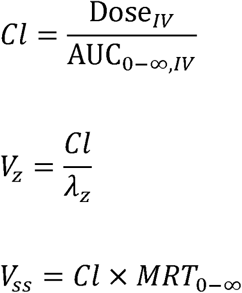

where *MRT_O—∞_*is the mean residence time extrapolated to infinity and was calculated as:

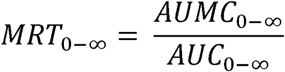

Where *AUMC_O—∞_*is the area under the (time x concentration) versus time curve extrapolated to infinity.

Following oral administration, systemic jugular-vein concentrations were used to estimate *C_max_*, *T_max_*, *y_z_*, *t_1/2_*, *AUC_O—t_*, and *AUC_O—∞_*. Apparent oral clearance (*Ci/F*) and apparent terminal-phase volume of distribution (*Y_z_/F*) were calculated as:

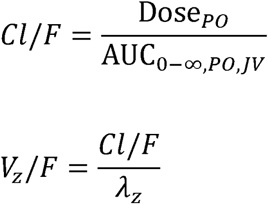

Absolute oral bioavailability (*F*) was estimated from dose-normalized systemic plasma exposure measured in the jugular vein following oral and IV administration:

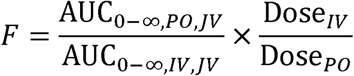

where *AUC_O—∞,Po,JV_* and *AUC_O—∞,fV,JV_* represent systemic jugular-vein exposure following oral and IV administration, respectively. Because the IV and oral studies were conducted in different animals, absolute bioavailability was estimated using the ratio of the corresponding treatment-group mean AUC values.

In addition to standard noncompartmental parameters, exploratory analyses were conducted to estimate hepatic extraction, hepatic availability, and the combined fraction of orally administered *ent*-verticilide absorbed and escaping intestinal first-pass loss. For exploratory calculations, systemic clearance following IV administration was assumed to predominantly represent hepatic clearance, and renal, biliary, and other nonhepatic clearance pathways were considered negligible. The blood-to-plasma concentration ratio of *ent*-verticilide was previously estimated to be approximately 0.85, indicating limited partitioning into erythrocytes relative to plasma. Because this ratio was considered sufficiently close to unity for the exploratory analysis, no correction between plasma and whole-blood clearance was applied.

Estimated hepatic extraction following IV administration was calculated as:

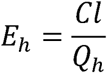

where *Ci* is systemic plasma clearance estimated by noncompartmental analysis following IV administration and rat hepatic blood flow (*Q_h_*) was assumed to be 70 mL/min/kg. The corresponding fraction escaping hepatic first-pass extraction was calculated as:

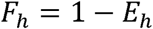

Because hepatic blood flow and nonhepatic clearance were not measured directly, the resulting *E_h_*and *F_h_* values were considered exploratory estimates rather than definitive hepatic pharmacokinetic parameters.

Mechanistically, absolute oral bioavailability was considered the product of the fraction absorbed from the intestinal lumen (*F_a_*), the fraction escaping intestinal metabolism and efflux (*F_g_*), and the fraction escaping hepatic first-pass extraction (*F_h_*):^16^

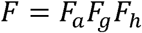

Accordingly, the combined fraction absorbed and escaping intestinal first-pass loss (intestinal availability) was estimated indirectly from absolute oral bioavailability and the IV-derived estimate of hepatic availability:

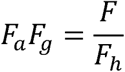

This calculation was considered exploratory because it depended on the assumptions that systemic clearance following IV administration predominantly represented hepatic clearance, nonhepatic clearance was negligible, hepatic blood flow was adequately represented by the assumed literature value, and pharmacokinetics were linear between the IV and oral dose levels.

As a separate exploratory approach to estimate intestinal (*F_a_F_g_*) and hepatic (*F_h_*) availability, *ent*-verticilide concentrations were measured in portal- and jugular-vein plasma following oral administration. Portal-vein *C_max_*, *T_max_*, *AUC_O—t_*, and *AUC_O—e,Po,PV_*were determined by noncompartmental analysis using the same procedures applied to the jugular-vein concentration–time profiles. Portal-vein concentrations following oral administration contain both newly absorbed drug and drug returning to the gastrointestinal circulation following systemic distribution.^17^ Therefore, total portal-vein AUC was not considered a direct measure of intestinal absorption. Net portal exposure attributable to intestinal absorption was estimated from the difference between portal-and jugular-vein exposure:^17^

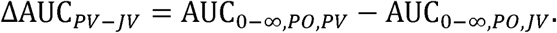

Under the assumptions that jugular-vein concentrations approximated the arterial concentrations entering the gastrointestinal circulation and that rat portal blood flow (*Q_PV_*) remained constant at 32.9 mL/min/kg,^17^ the portal–systemic estimate of *F_a_F_g_*was calculated as:^17^

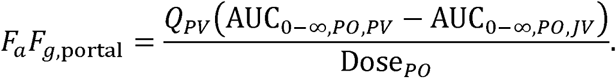

Because the blood-to-plasma concentration ratio was considered sufficiently close to unity, no additional blood-to-plasma correction was applied. The portal-derived estimate was compared descriptively with the indirect *F_a_F_g_* estimate derived from *F/F_h_*. The portal-derived calculation was considered exploratory because portal blood flow was not measured directly, jugular-vein concentrations were used as a surrogate for arterial concentrations supplying the gastrointestinal tract, and cannulation, formulation, blood sampling, or other study procedures may have altered portal blood flow. The apparent *F_h_* was calculated as:

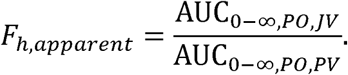

Because jugular-vein concentrations do not represent direct hepatic venous outflow, portal-vein exposure includes systemically recirculating drug, and hepatic arterial input was not measured, *F_h,opporent_* was considered exploratory indices of hepatic availability rather than definitive estimates.

The effects of ABT pretreatment were evaluated by comparing portal- and jugular-vein concentration–time profiles and pharmacokinetic parameters between vehicle- and ABT-pretreated animals. Comparisons included *C_max_*, *T_max_*, *t_1/2_*, *AUC_O—t_*, *AUC_O—e_*, *CL/F*, *Y_z_/F*, relative exposure ratios, and the portal–systemic AUC difference. Because ABT inhibits CYP activity in both the intestine and liver and may also reduce systemic metabolic clearance after absorption, changes in systemic exposure following ABT pretreatment were interpreted as evidence for a contribution of CYP-mediated metabolism but were not considered sufficient to localize metabolism specifically to the intestinal wall or liver. Portal and systemic exposure patterns were evaluated descriptively to explore the potential relative contributions of intestinal and hepatic metabolism.

Pharmacokinetic parameters were calculated for individual animals and summarized by treatment groups. Continuous pharmacokinetic parameters were summarized using the arithmetic mean and standard deviation. Parameters derived from ratios of treatment-group mean values, including absolute oral bioavailability and the indirect estimate of *F_a_F_g_*, were identified as group-level estimates rather than means of individual-animal parameters.

### LC–MS/MS Bioanalysis

#### Analysis of In Vitro Study Samples

Samples generated from the TRANSIL protein-binding, biorelevant-media stability, hepatic microsomal stability, and Caco-2 permeability experiments were analyzed by liquid chromatography–tandem mass spectrometry (LC–MS/MS) using multiple-reaction monitoring. Samples were introduced using a CTC Analytics PAL-xt DW autosampler and separated using a Shimadzu LC-20AD Series HPLC system coupled to a SCIEX 6500 QTRAP mass spectrometer (SCIEX, Foster City, CA, USA). Injection volumes were 5 µL for the TRANSIL and biorelevant-media stability assays and 10 µL for the hepatic microsomal stability and Caco-2 permeability assays. The mass spectrometer was equipped with an electrospray ionization source operated in positive-ion mode with an ion-source temperature of 500 °C and an ion-spray voltage of 4,500 V. Chromatographic separation was performed using a Kinetex C18 column (50 × 2.1 mm, 1.7-µm particle size; Phenomenex, Torrance, CA, USA) maintained at 60 °C. Mobile phase A consisted of 0.1% formic acid in ultrapure water, and mobile phase B consisted of 0.1% formic acid in acetonitrile. The mobile phases were delivered at a flow rate of 0.5 mL/min. The initial mobile-phase composition was 5% B. Mobile phase B was increased linearly from 5% to 90% over 0.5 min and maintained at 90% from 0.5 to 2.0 min. The composition was returned to 5% B between 2.00 and 2.01 min, and the column was re-equilibrated at 5% B through the end of the 2.5-min analytical run. *ent*-Verticilide was monitored using the precursor-to-product-ion transition *m/z* 853.6→214.1. Verapamil and carbamazepine were monitored using the transitions 455.0→165.0, and 237.1→194.0, respectively. Chromatographic peak areas were integrated using Analyst software version 1.7.0.

For the TRANSIL binding assays, the integrated *ent*-verticilide peak area was used as the analytical response after LC–MS/MS response linearity was confirmed across the evaluated concentration range. Responses from wells containing immobilized protein were compared with responses from reference wells without immobilized protein to account for nonspecific binding. Binding parameters were calculated using the manufacturer-provided TRANSIL analysis procedure.

For the biorelevant-media and hepatic microsomal stability assays, the analytical response was calculated as:

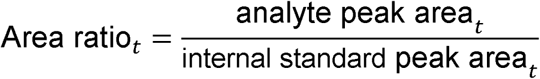

The percentage of *ent*-verticilide remaining at each time point was calculated relative to the response at time zero:

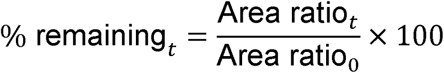

For the hepatic microsomal stability experiments, carbamazepine was used as the internal standard. The percentages of *ent*-verticilide and the positive-control substrate verapamil remaining at each time point were calculated from their respective analyte-to-carbamazepine peak-area ratios relative to the corresponding time-zero ratios.

### Quantification of Caco-2 Permeability Samples

Matrix-matched calibration standards were prepared separately in Caco-2 assay matrices with and without bovine serum albumin to produce final *ent*-verticilide concentrations ranging from 20 to 20,000 nM. Quality control (QC) samples were prepared independently in the corresponding matrices at nominal concentrations of 120, 600, 3,000, and 15,000 nM. A 25-µL aliquot of each calibration standard, QC sample, or Caco-2 study sample was transferred to a 96-well plate containing 75 µL of acetonitrile with internal standard. The resulting extracts were diluted twofold with ultrapure water, and 10 µL was injected for LC–MS/MS analysis using the chromatographic and mass-spectrometric conditions described above. *ent*-Verticilide was monitored using the transition *m/z* 853.6→214.1. Atenolol, metoprolol, digoxin, and tolbutamide were monitored using the transitions *m/z* 267.2→145.0, 268.2→148.0, 798.5→651.5, and 271.1→155.0, respectively. Tolbutamide served as the internal standard. Calibration curves were constructed by plotting the *ent*-verticilide-to-internal-standard peak-area ratio against nominal *ent*-verticilide concentration. Study-sample concentrations were determined using weighted quadratic regression with a weighting factor of 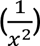. An analytical run was accepted when the calibration curve contained at least five standards with back-calculated concentrations within ±15% of their nominal concentrations and a coefficient of determination (*R*^2^) greater than 0.90. At least 66.7% of all QC samples and at least 50% of the QC samples at each concentration were required to be within ±15% of their nominal concentrations.

### Quantification of Rat Pharmacokinetic Samples

Matrix-matched calibration standards were prepared by adding 5 µL of an *ent*-verticilide working solution to 50 µL of blank male Sprague–Dawley rat plasma to produce final concentrations ranging from 0.5 to 1,000 ng/mL. QC samples were prepared independently at nominal concentrations of 1, 2, 50, and 800 ng/mL. Calibration standards, QC samples, and plasma samples collected during the rat pharmacokinetic studies were thawed at room temperature. A 55-µL aliquot of each sample was transferred to a protein-precipitation plate containing 200 µL of ice-cold acetonitrile containing internal standard. Carbamazepine was used as the internal standard for samples from the IV/PO pharmacokinetic study, whereas verapamil was used for samples from the PO/ABT pharmacokinetic study. Samples were vortex-mixed for 30 s and centrifuged at 4,000 rpm for 15 min at 4 °C. The resulting supernatant was diluted threefold with ultrapure water, and 12 µL was injected for LC–MS/MS analysis.

Chromatographic separation was performed using a Shimadzu LC-30AD Series HPLC system equipped with a SIL-30AC autosampler (Shimadzu Corporation, Kyoto, Japan). The HPLC system was coupled to an AB SCIEX API 4000 mass spectrometer for analysis of samples from the IV/PO study and a SCIEX 5500 mass spectrometer for analysis of samples from the PO/ABT study (SCIEX, Foster City, CA, USA). Both instruments were operated using electrospray ionization in positive-ion mode. Analytes were separated on a HALO AQ-C18 column (30 × 3.0 mm, 2-µm particle size; Advanced Materials Technology, Wilmington, DE, USA). Mobile phase A consisted of 0.1% formic acid in ultrapure water, and mobile phase B consisted of 0.1% formic acid in acetonitrile–water (95:5, v/v). The mobile phases were delivered at a flow rate of 0.6 mL/min. The initial composition of 30% B was maintained through 0.2 min. Mobile phase B was increased linearly from 30% to 100% between 0.2 and 1.6 min and maintained at 100% through 2.30 min. The composition was returned to 30% B between 2.31 and 2.50 min, and the column was re-equilibrated at the initial composition before the subsequent injection. *ent*-Verticilide was monitored using the precursor-to-product-ion transition *m/z* 853.6→214.1. Carbamazepine and verapamil were monitored using the transitions *m/z* 237.0→194.2 and 455.3→165.2, respectively.

Calibration curves were constructed by plotting the *ent*-verticilide-to-internal-standard peak-area ratio against nominal *ent*-verticilide concentration. Study-sample concentrations were determined using weighted linear regression with a weighting factor of 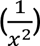. An analytical run was accepted when the calibration curve contained at least five standards with back-calculated concentrations within ±15% of their nominal concentrations and an (*R*^2^) greater than 0.90. At least 66.7% of all QC samples and at least 50% of the QC samples at each concentration were required to be within ±15% of their nominal concentrations.

## RESULTS

### Stability and Nonspecific Binding in Biorelevant Media

*ent*-Verticilide concentrations remained stable over the 120-minute incubation period in rat plasma and FaSSIF at pH 6.5 (**Fig 1, A**). Mean peak-area ratios in these matrices generally remained near or above the baseline value throughout the experiment, with approximately 103–113% of baseline detected at 120 minutes (**Fig. 1, A**). These findings indicated no measurable loss of *ent*-verticilide under the evaluated plasma or simulated intestinal conditions.

**Fig. 1:**
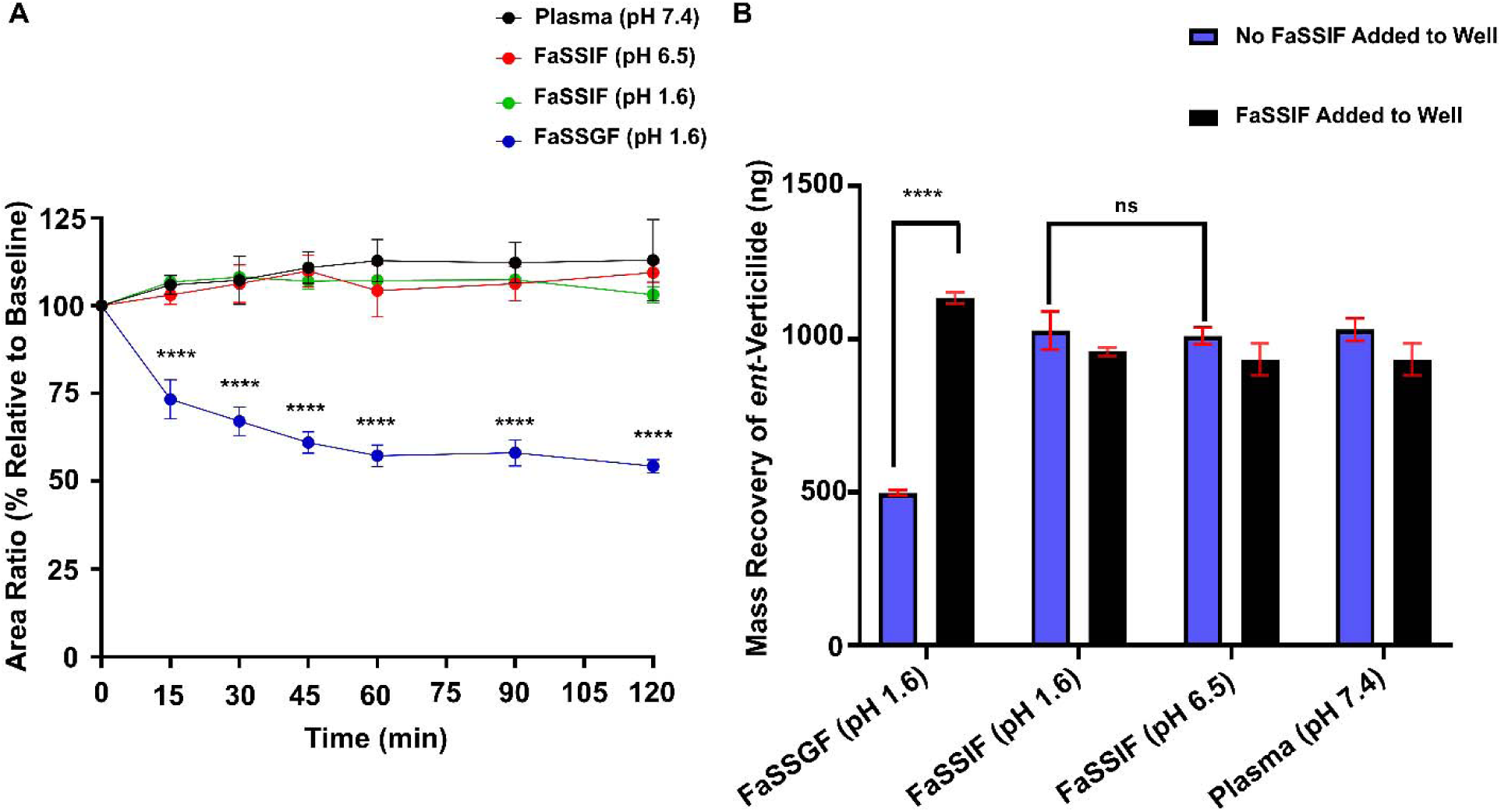
In Vitro Stability (**A**) and Nonspecific Binding (**B**) of ent-Verticilide in Biorelevant Media. Data were shown as mean ± S.D. For panel (**A**), n = 3–6 for each concentration. **** indicates statistical significance between FaSSGF (pH 1.6) and the other conditions at each time point (p < 0.0001), as determined by two-way ANOVA with Dunnett’s correction for multiple comparisons. For panel (**B**), n = 3 for each condition and only relevant comparisons were made and shown in the figure. ****p < 0.0001 by two-way ANOVA with Dunnett’s correction for multiple comparisons. **ns**: Not significant.

In contrast, the measured concentration of *ent*-verticilide declined rapidly in FaSSGF at pH 1.6 (**Fig. 1, A**). Approximately 73% of the baseline peak-area ratio remained at 15 minutes, followed by a more gradual decline to approximately 54% at 120 minutes (**Fig. 1, A**). Because *ent*-verticilide remained stable in acidified FaSSIF (pH 1.6) and in FaSSIF (pH 6.5) (**Fig. 1, A**), the apparent loss in FaSSGF could not be attributed to acidic conditions alone.

The addition of fresh FaSSIF to the wells after the stability experiment substantially increased the recovery of *ent*-verticilide from wells containing FaSSGF. Before FaSSIF addition, approximately 500 ng of *ent*-verticilide was recovered from the FaSSGF (pH 1.6) wells (**Fig. 1, B**). Following the addition of FaSSIF and further incubation, the recovered mass increased to approximately 1,130 ng (**Fig. 1, B**), corresponding to more than a two-fold increase.

In contrast, FaSSIF addition did not increase recovery from wells containing acidified FaSSIF, FaSSIF at pH 6.5, or plasma (**Fig. 1, B**). Recovery before FaSSIF addition was approximately 1,000–1,030 ng in these matrices and remained approximately 930–960 ng after the additional incubation. The selective recovery of *ent*-verticilide from the FaSSGF wells indicates that its initial disappearance from the aqueous phase represented reversible nonspecific binding to the sample wells rather than degradation.

### In Vitro Plasma Protein Binding Assay

*ent*-Verticilide exhibited extensive plasma protein binding across the concentration range of 0.01––10 µM (**Table 1**). In the PPB assay, *ent*-verticilide was 99.1%, 99.5%, 98.4%, and 94.2% bound at 0.01, 0.1, 1, and 10 µM, respectively. The corresponding unbound fractions were 0.9%, 0.5%, 1.6%, and 5.8%. In the human serum albumin assay, *ent*-verticilide was 96.4%, 96.9%, 95.9%, and 92.7% bound at the respective concentrations, corresponding to unbound fractions of 3.6%, 3.1%, 4.1%, and 7.3%. Warfarin, included as a high-protein-binding reference compound at 1 µM, was 97.1% bound in the PPB assay and 98.4% bound in the HSA assay. Collectively, these results demonstrated that *ent*-verticilide was extensively bound to plasma proteins, with albumin representing a major binding component. Binding remained greater than 98% in the plasma protein-binding assay at concentrations up to 1 µM, with a modest reduction in binding observed at 10 µM.

### In Vitro Permeability Assay

The performance of the Caco-2 system was evaluated using established reference compounds. Atenolol exhibited low A-to-B permeability, with a *P_app_*of 0.460 x 10^−6^ cm/s, whereas metoprolol exhibited substantially higher A-to-B permeability of 18.050 x 10^−6^ cm/s (**Table 2**). Digoxin demonstrated directional transport, with A-to-B and B-to-A *P_app_*values of 1.585 x 10^−6^ and 12.200 x 10^−6^ cm/s, respectively, and an efflux ratio of 7.70 (**Table 2**). These results confirmed the ability of the assay to distinguish compounds with low and high passive permeability and to detect transporter-associated efflux.

**Table 2:** In Vitro Permeability of *ent*-Verticilide in Caco-2 Cells. Atenolol, Metoprolol, and Digoxin were included as reference compounds. ***ent*-vert**: *ent*-Verticilide; **BSA**: Bovine serum albumin. *P_app_*: Apparent permeability; *P_gp_*: P-glycoprotein

| Compound | Caco-2<br>Assay<br>Control | Concentration<br>($\mu\text{M}$) | $P_{app}$ ($\times 10^{-6}$ cm/s) | | Efflux Ratio<br>(B-A/A-B) | % Recovery | |
| --- | --- | --- | --- | --- | --- | --- | --- |
|  |  |  | A-B | B-A |  | A-B | B-A |
| Atenolol | Low | 10 | 0.460 | 1.409 | 3.06 | 105 | 109 |
| Metoprolol | High | 10 | 18.050 | 15.475 | 0.86 | 105 | 99 |
| Digoxin | $P_{gp}$<br>Substrate | 10 | 1.585 | 12.200 | 7.70 | 94 | 92 |
| *ent*-vert | N/A | 10 | 0.895 | 1.250 | 1.40 | 25 | 84 |
| *ent*-vert + BSA | N/A | 10 | 1.420 | 1.820 | 1.28 | 48 | 123 |

*ent*-Verticilide exhibited low bidirectional permeability. In the absence of bovine serum albumin (BSA), its A-to-B and B-to-A *P_app_*values were 0.895 x 10^−6^ and 1.250 x 10^−6^ cm/s, respectively (**Table 2**). The A-to-B permeability of *ent*-verticilide was approximately two-fold greater than that of the low-permeability control atenolol but approximately 21-fold lower than that of the high-permeability control metoprolol. The addition of BSA modestly increased the A-to-B and B-to-A *P_app_* values to 1.420 x 10^−6^ and 1.820 x 10^−6^ cm/s, respectively (**Table 2**). Efflux ratios remained below 2 both without and with BSA (1.40 and 1.28, respectively), providing no evidence of pronounced net efflux under the evaluated conditions.

Mass recovery of the reference compounds ranged from 92% to 109% (**Table 2**). In contrast, *ent*-verticilide recovery was direction dependent and substantially lower, particularly in the A-to-B experiment, in which only 25% was recovered. Recovery in the B-to-A experiment was 84%. The addition of BSA increased recovery to 48% in the A-to-B direction and 123% in the B-to-A direction. Although BSA improved recovery, acceptable mass balance was not achieved consistently.

### Metabolic Stability in Rat and Human Liver Microsomes

The metabolic stability of *ent*-verticilide was evaluated in rat and human liver microsomes in the presence and absence of NADPH (**Fig. 2**). Verapamil, a known CYP3A substrate, was included as a positive control to confirm the metabolic activity of the microsomal fractions (data not shown). In the absence of NADPH, *ent*-verticilide remained generally stable in both species, with approximately 80–100% of the initial concentration remaining throughout the 45-min incubation. In contrast, addition of NADPH resulted in rapid disappearance of *ent*-verticilide. At 15 min, approximately 9% and 14% of the initial concentration remained in human and rat liver microsomal incubations, respectively. The pronounced difference between incubations with and without NADPH demonstrated that microsomal disappearance of *ent*-verticilide was predominantly NADPH dependent; this finding is consistent with metabolism mediated primarily by CYP enzymes.

**Fig. 2.**
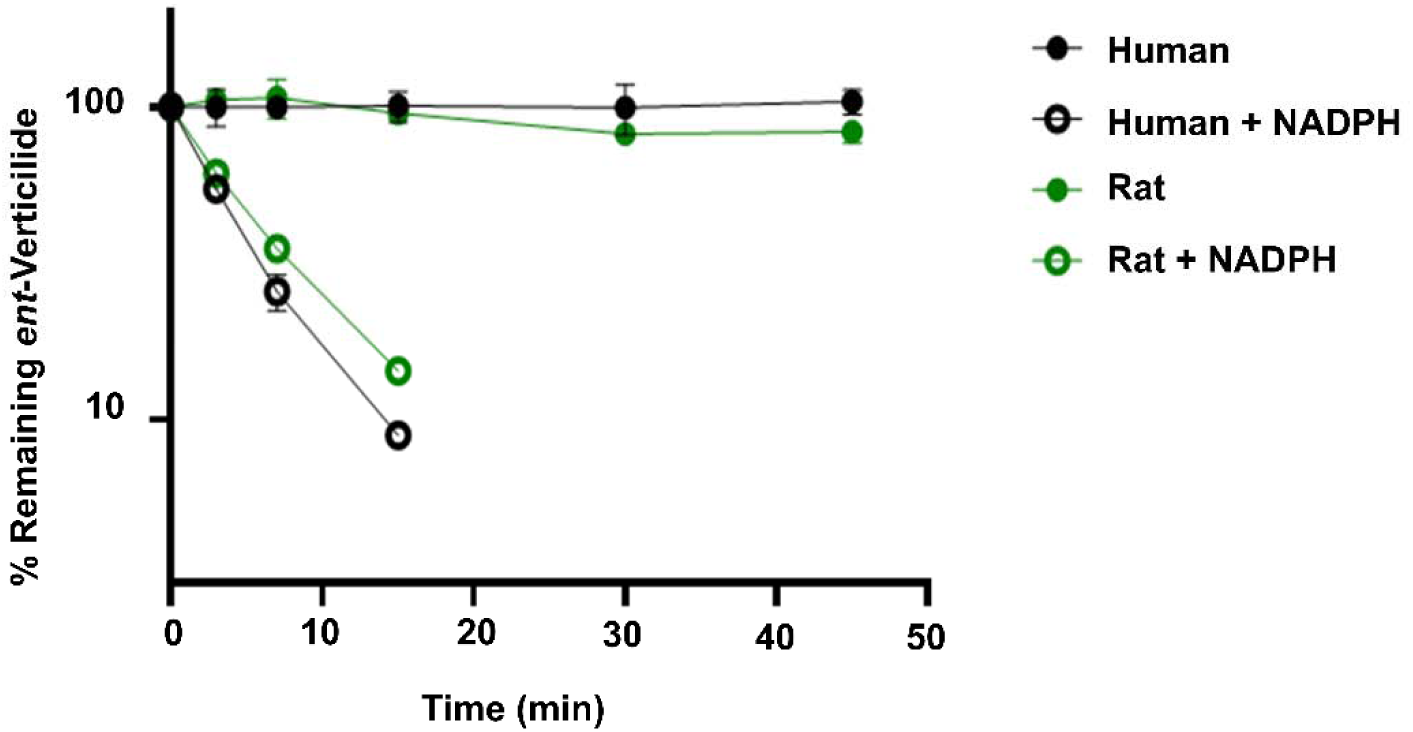
*ent*-Verticilide Undergoes NADPH-Dependent Metabolism in Rat and Human Hepatic Microsomes. *ent*-Verticilide was incubated with rat or human liver microsomes in the presence or absence of NADPH. The percentage remaining at each time point was calculated from the *ent*-verticilide-to-internal-standard peak-area ratio relative to the corresponding time-zero ratio. Data were presented as mean ± S.D from three replicate incubations per time point. The percentage remaining is displayed on a logarithmic scale, and the log-transformed data were fitted using simple linear regression. **NADPH**: nicotinamide adenine dinucleotide phosphate.

Scaling of the microsomal disappearance data produced estimated intrinsic clearance values of 748 and 255 mL/min/kg in rat and human, respectively (**Table 3**). Application of the binding-unadjusted well-stirred hepatic model resulted in predicted hepatic clearances of 64.0 mL/min/kg in rat and 19.4 mL/min/kg in human. The corresponding predicted hepatic extraction ratios were 0.91 and 0.92, respectively, indicating high predicted hepatic extraction in both species under the assumptions of the model.

**Table 3:** Predicted Parameters from Substrate Depletion of *ent*-Verticilide in Rat and Human Hepatic Microsomes. *k*: First-order elimination rate constant; *t_1/2_*: In vitro half-life; *cL_int,app,scaled_*: Apparent hepatic intrinsic clearance, scaled; *cL_n_*: Predicted hepatic clearance; *z_n_*: Predicted hepatic extraction ratio.

| Species | $k$<br>(min<sup>-1</sup>) | $t_{1/2}$<br>(min) | $CL_{int,app,scaled}$<br>(mL/min/kg) | $CL_H$<br>(mL/min/kg) | $E_H$ |
| --- | --- | --- | --- | --- | --- |
| Rat | 0.13 | 5.43 | 623 | 63 | 0.90 |
| Human | 0.16 | 4.35 | 255 | 19.4 | 0.92 |

### In Vivo Pharmacokinetics in Rats

#### Pharmacokinetics Following Intravenous and Oral Administration

The mean plasma concentration-time profile following a single 1 mg/kg IV dose is presented in **Fig. 3**, and the corresponding pharmacokinetic parameters are summarized in **Table 4**. The estimated concentration at time zero was 1,220 ± 857 ng/mL and rapidly declined during the initial distribution phase, followed by a slower terminal phase. The terminal elimination rate constant was 0.134 ± 0.008 *hr*^−1^, corresponding to a terminal half-life of 5.19 ± 0.30 h. The mean *AUC_O—∞_* was 670 ± 131 ng·h/mL with a systemic clearance of 25.5 ± 4.5 mL/min/kg. The multiexponential decline in plasma concentrations with a time-dependent increase in volume of distribution from 1.10 ± 0.61 L/kg (*Y_c_*), 4.72 ± 1.44 L/kg (*Y_ss_*), and 11.4 ± 2.0 L/kg (*Y_z_*) is consistent with multicompartment disposition and extensive distribution of *ent*-verticilide beyond the central plasma compartment.

**Fig. 3.**
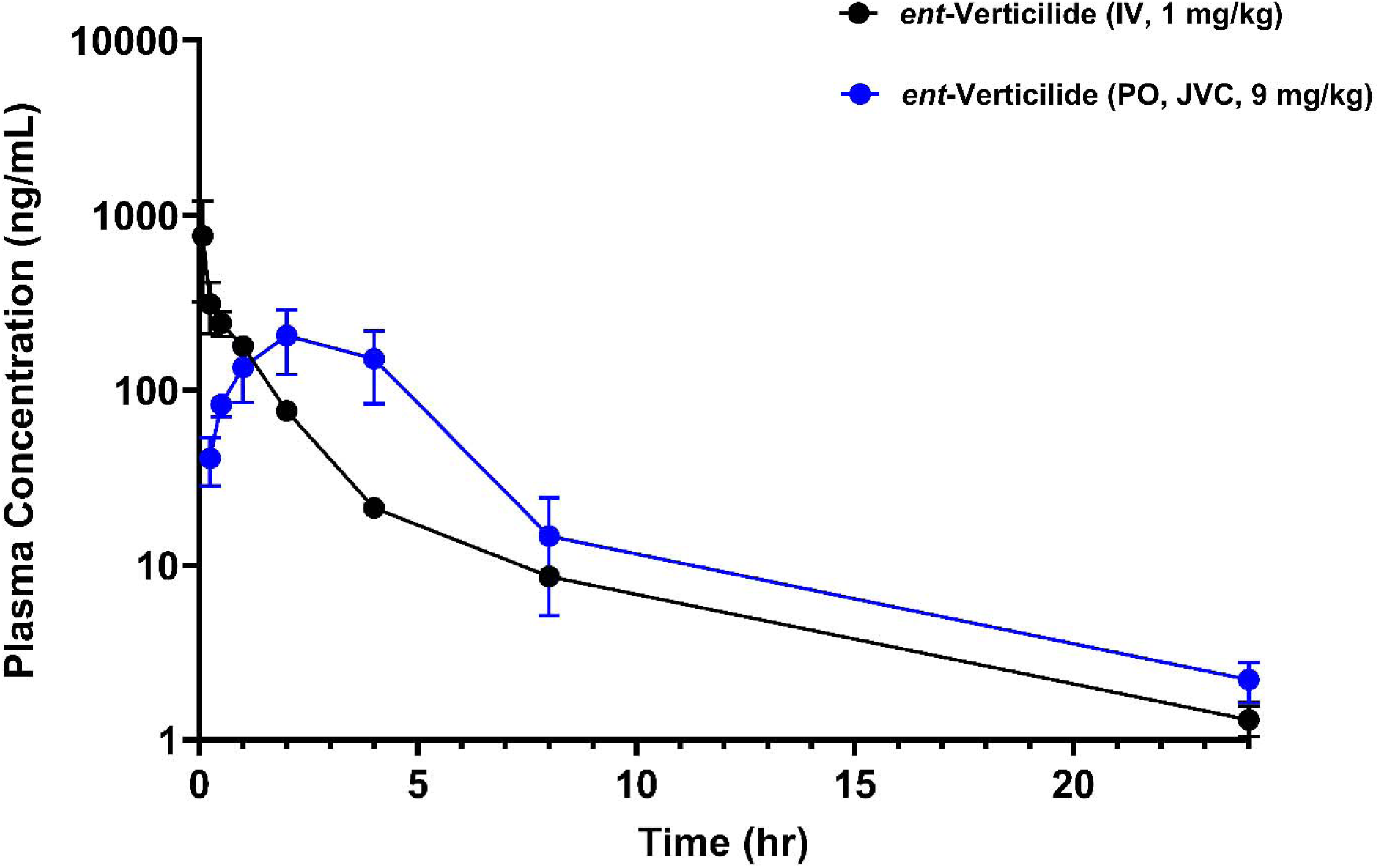
Plasma-Concentration-versus-Time Profile of *ent*-Verticilide Following IV (1 mg/kg) and PO (9 mg/kg) Administration in Rat. For PO route, blood samples were collected via jugular vein cannulation (JVC). Data = mean ± S.D. from n = 3 for each concentration.

**Table 4:** Non-Compartmental Analysis of Pharmacokinetic Parameters of *ent*-Verticilide following PO and IV Administration in Rat. For oral administration, blood samples were collected via jugular vein (JVC) for assessment of systemic exposure. Bioavailability (F) for IV route was assumed to be one. Data were shown as mean ± S.D. *k_e_*: In vivo elimination rate constant; *T_1/2_*: In vivo half-life; *c_o_*: Initial plasma concentration; *c_max_*: Maximum observed plasma concentration; *T_max_*: Time to *c_max_*; *Kvc_o—∞_*: Area under the concentration-versus-time curve from time 0 to infinity; *cL*: Observed clearance; *cL/F*: apparent oral clearance; *r_c_*: Volume of distribution of the central compartment; *r_ss_*: Volume of distribution at steady state; *r_z_*: Terminal volume of distribution; *r_z_/F*: Apparent terminal volume of distribution; *F*: Absolute oral bioavailability. **N/A**: Not applicable

| IV Parameters | Mean ± S.D | PO Parameters | Mean ± S.D |
| --- | --- | --- | --- |
| Dose (mg/kg) | 1.0 | Dose (mg/kg) | 9.0 |
| $k_e$ ($hr^{-1}$) | $0.134 \pm 0.008$ | $k_e$ ($hr^{-1}$) | $0.181 \pm 0.019$ |
| $T_{1/2}$ (hr) | $5.19 \pm 0.30$ | $T_{1/2}$ (hr) | $3.85 \pm 0.38$ |
| $C_0$ (ng/mL) | $1220 \pm 857$ | $C_{max}$ (ng/mL) | $205 \pm 82$ |
| $T_{max}$ (hr) | N/A | $T_{max}$ (hr) | $2.00 \pm 0.00$ |
| $AUC_{0-\infty}$ (ng·h/mL) | $670 \pm 131$ | $AUC_{0-\infty}$ (ng·h/mL) | $946 \pm 338$ |
| $CL$ (mL/min/kg) | $25.5 \pm 4.5$ | $CL/F$ (mL/min/kg) | $176 \pm 72.5$ |
| $V_c$ (L/kg) | $1.10 \pm 0.61$ | $V_c$ (L/kg) | N/A |
| $V_{ss}$ (L/kg) | $4.72 \pm 1.44$ | $V_{ss}$ (L/kg) | N/A |
| $V_z$ (L/kg) | $11.4 \pm 2.0$ | $V_z/F$ (L/kg) | $59.3 \pm 26.9$ |
| $F$ | 1.00 | $F$ | $0.16 \pm 0.06$ |

The mean plasma concentration-time profile following a single 9 mg/kg PO dose is presented in **Fig. 3**, and the corresponding pharmacokinetic parameters are summarized in **Table 4**. Following oral administration, *ent*-verticilide reached a maximum jugular-vein plasma concentration of 205 ± 82 ng/mL with a *T_mox_* of 2.00 hours. Plasma concentrations subsequently declined, with quantifiable concentrations remaining through 24 h. The mean oral *AUC_O—∞_* was 946 ± 338 ng·h/mL, and the terminal elimination half-life was 3.85 ± 0.38 h. Apparent oral clearance and terminal-phase volume of distribution were 176 ± 72.5 mL/min/kg and 59.3 ± 26.9 L/kg, respectively. Dose-normalized systemic exposure following oral and intravenous administration yielded a mean absolute oral bioavailability of *ent*-verticilide of 0.16 ± 0.06, indicating that 16% of the administered oral dose reached the systemic circulation.

### Portal and Jugular Pharmacokinetics With and Without ABT Pretreatment

To explore the intestinal availability of *ent*-verticilide, a single oral dose of 9 mg/kg was administered to rats with portal- and jugular-vein cannulas following pretreatment with ABT or vehicle. Serial blood sampling from both vascular compartments permitted comparison of portal and systemic concentration–time profiles within each treatment group. The mean concentration–time profiles are presented in **Fig. 4**, and the corresponding noncompartmental pharmacokinetic parameters are summarized in **Table 5**.

**Fig. 4:**
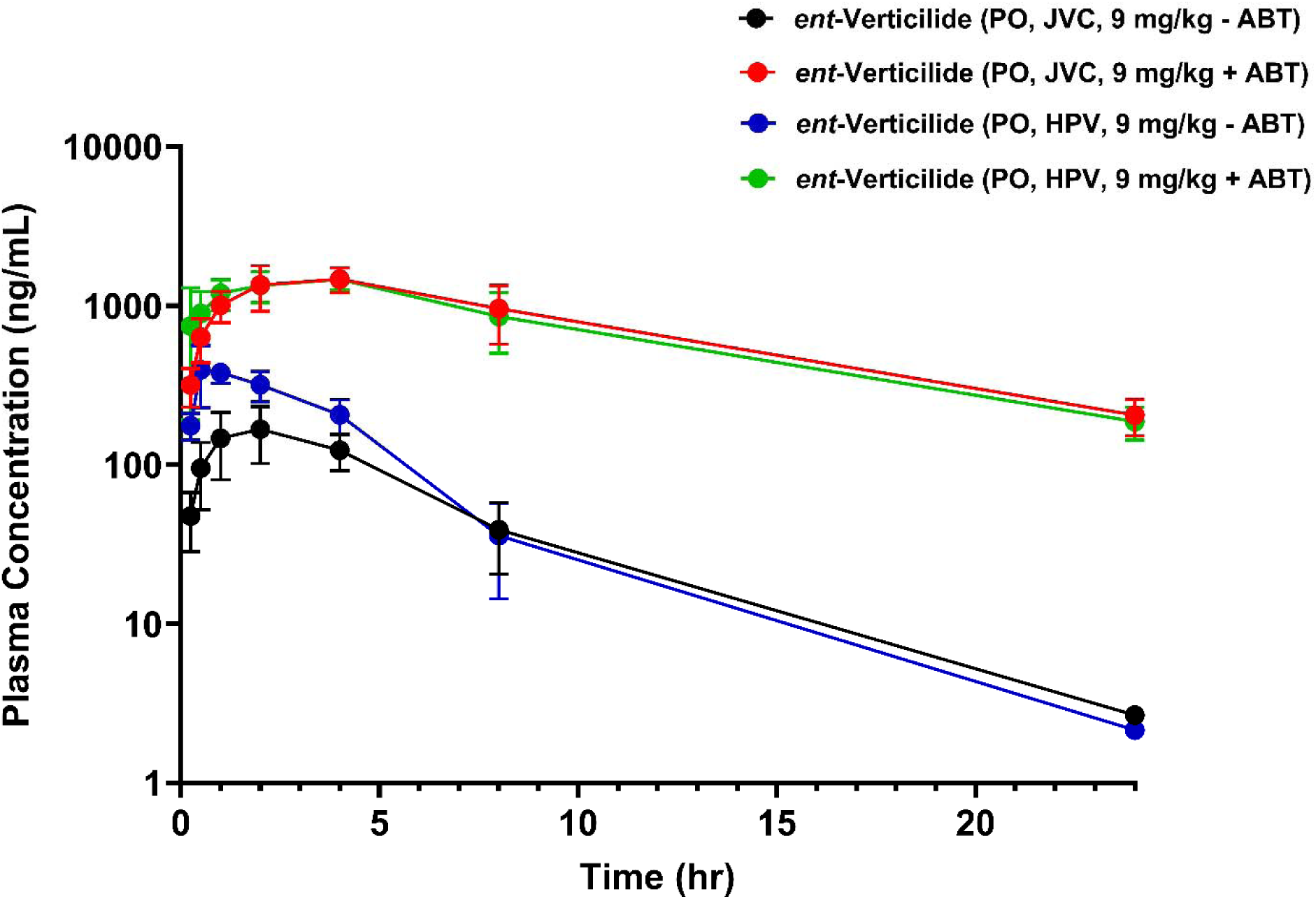
Plasma-Concentration-versus-Time Profile of *ent*-Verticilide Following PO (9 mg/kg) Administration in Rat. Blood samples were collected via jugular vein cannulation (JVC) and hepatic portal vein (HPV), with and without the presence of the CYP inhibitor 1-aminobenzotriazole (ABT). Data = mean ± S.D. from n = 3 for each concentration.

**Table 5:**
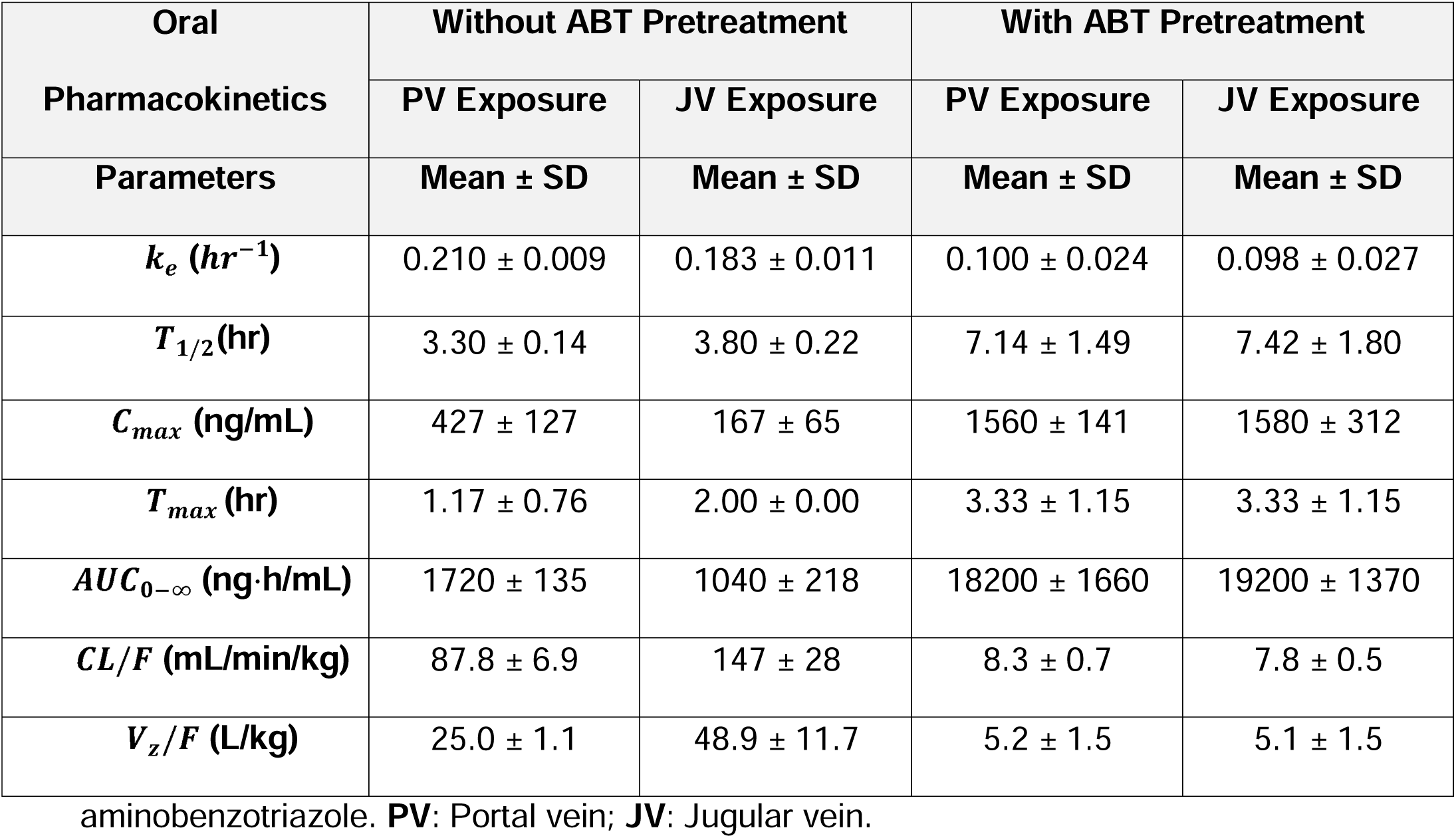
Non-Compartmental Analysis of Pharmacokinetic Parameters of *ent*-Verticilide following PO (9 mg/kg) Administration in Rat, in the Presence or Absence of CYP Inhibitor ABT (100 mg/kg). *ent*-Verticilide was administered orally following pretreatment with ABT (100 mg/kg) or vehicle. Pharmacokinetic parameters were determined from portal- and jugular-vein plasma concentration–time profiles. Data are presented as the mean ± SD for three animals per treatment group. **ABT**: 1-

In rats not pretreated with ABT, portal-vein concentrations exceeded jugular-vein concentrations during the absorption phase. Portal *C_mox_* was 427 ± 127 ng/mL compared with a jugular *C_mox_* of 167 ± 65 ng/mL, representing an approximately 2.6-fold portal-to-jugular difference. Portal *AUC_O—∞_* was 1,720 ± 135 ng·h/mL compared with 1,040 ± 218 ng·h/mL in jugular plasma, representing an approximately 1.7-fold difference. *T_mox_* occurred earlier in portal plasma than in jugular plasma, with values of 1.17 and 2.00 h, respectively. Terminal half-lives were similar between sampling compartments, with values of 3.30 ± 0.14 h in portal plasma and 3.80 ± 0.22 h in jugular plasma.

ABT pretreatment markedly increased *ent*-verticilide concentrations in both portal and jugular plasma. Portal *C_mox_* increased approximately 3.7-fold, from 427 ± 127 to 1,560 ± 141 ng/mL, whereas jugular *C_mox_* increased approximately 9.5-fold, from 167 ± 65 to 1,580 ± 312 ng/mL. Portal *AUC_O—∞_* increased approximately 10.6-fold, from 1,720 ± 135 to 18,200 ± 1,660 ng·h/mL. Jugular *AUC_O—∞_* increased approximately 18.5-fold, from 1,040 ± 218 to 19,200 ± 1,370 ng·h/mL.

ABT pretreatment also prolonged the terminal elimination phase. The terminal half-life increased from 3.30 ± 0.14 to 7.14 ± 1.49 h in portal plasma and from 3.80 ± 0.22 to 7.42 ± 1.80 h in jugular plasma. *T_mox_* was delayed to 3.33 ± 1.15 h in both compartments. Apparent oral clearance decreased from 87.8 ± 6.9 to 8.3 ± 0.7 mL/min/kg based on portal exposure and from 147 ± 28 to 7.8 ± 0.5 mL/min/kg based on jugular exposure. Following ABT pretreatment, the portal- and jugular-vein concentration–time profiles were nearly superimposable, with comparable *C_mox_*, *T_mox_*, *AUC_O—∞_*, and terminal half-life values.

### Exploratory Mechanistic Oral Bioavailability Analysis

Exploratory pharmacokinetic calculations were performed to assess the potential contributions of hepatic extraction and processes occurring before hepatic entry to the low oral bioavailability of *ent*-verticilide. Systemic clearance following intravenous administration was assumed to predominantly represent hepatic clearance, and rat hepatic blood flow was assumed to be 70 mL/min/kg. Under these assumptions, the fraction escaping hepatic extraction *F_h_* was estimated to be 0.64 ± 0.06. Combining the estimated *F_h_* with the measured absolute *F* of 0.16 ± 0.06 produced an estimated *F_a_F_g_* of 0.25 ± 0.02.

In the portal–jugular study, an independent *F_a_F_g_*estimate was calculated from the difference between portal- and jugular-vein *AUC_O—∞_* following oral administration without ABT. This approach produced an *F_a_F_g_* estimate of 0.15 ± 0.07. The corresponding exploratory estimates of *F_h_*and overall *F* were calculated to be 0.62 ± 0.17 and 0.08 ± 0.03, respectively. Following ABT pretreatment, portal *AUC_O—∞_* was not greater than jugular *AUC_O—∞_*. Consequently, subtraction of jugular exposure from portal exposure did not produce a physiologically meaningful estimate of net intestinal input, and *F_a_F_g_*was not calculated for the ABT-treated group.

## DISCUSSION

The present study characterized the biopharmaceutical and pharmacokinetic properties of *ent*-verticilide and identified the principal factors governing its systemic exposure. Importantly, *ent*-verticilide was stable in rat plasma and simulated gastrointestinal fluids, indicating that chemical or enzymatic degradation in these matrices is unlikely to represent a major barrier to systemic exposure. This stability may reflect its D-alanine residues and the conformational constraint imposed by its cyclic depsipeptide backbone, which could reduce accessibility of its ester and amide linkages to hydrolytic enzymes. Although an apparent loss of *ent*-verticilide was observed in FaSSGF, this effect was largely reversed by the addition of FaSSIF, supporting reversible nonspecific binding rather than degradation as the underlying mechanism. The lower bile salt and phospholipid content of FaSSGF may provide insufficient solubilization and favor partitioning onto polypropylene surfaces, whereas the higher concentrations in FaSSIF likely enhance solubilization and recovery. Consistent with this interpretation, *ent*-verticilide exhibited extensive plasma protein binding, as commonly observed for lipophilic compounds,^18,19^ and plasma proteins likely improve in vitro recovery by maintaining the compound in solution and reducing adsorption to hydrophobic surfaces. Thus, while stability and nonspecific loss do not appear to be major intrinsic barriers to exposure, the permeability and metabolic data identify more consequential limitations to *ent*-verticilide disposition. Specifically, low apparent permeability and rapid NADPH-dependent hepatic metabolism emerged as the major liabilities likely restricting systemic exposure. Nevertheless, oral administration produced meaningful systemic exposure in rats, with an absolute bioavailability of approximately 16%. Moreover, the marked increases in both portal and systemic exposure following ABT pretreatment provide in vivo evidence that CYP-mediated metabolism contributes substantially to limiting *ent*-verticilide exposure and represents a potentially modifiable determinant of its pharmacokinetic profile.

In Caco-2 monolayers, *ent*-verticilide exhibited low absorptive apparent permeability, while efflux ratios below 2 provided no evidence for pronounced directional efflux under the conditions evaluated. Incomplete recovery, particularly in the absence of albumin, limits quantitative interpretation and is consistent with the nonspecific binding observed in other assay formats; thus, limited passive transcellular permeability, rather than active efflux, appears more likely to constrain intestinal absorption. Nevertheless, an absolute oral bioavailability of approximately 16% indicates that meaningful gastrointestinal absorption occurs despite the low apparent Caco-2 permeability. The cyclic depsipeptide structure of *ent*-verticilide may facilitate membrane transit by conformationally constraining the backbone and reducing its effective polarity, consistent with studies of *ent*-verticilide-inspired macrocycles showing that backbone composition can strongly influence passive permeability independently of side-chain composition.^20–22^ In addition to these backbone features, the pentyl side chain of *ent*-verticilide appears to play an important role in maintaining its passive permeability. In our previous study, replacement of a single pentyl side chain with a carboxylic acid-containing substituent resulted in a marked reduction in passive permeability,^23^ highlighting the importance of side-chain physicochemical properties in membrane transit. Together, these findings suggest that the permeability of *ent*-verticilide arises from a combination of favorable backbone conformation and side-chain hydrophobicity.

Cyclosporine A provides a complementary example of how large cyclic peptides can achieve meaningful oral exposure through a combination of favorable conformational properties, solubilization, transport, and metabolism.^6^ Carrier-mediated uptake has also been demonstrated for an engineered cyclosporine–thiodipeptide conjugate, while intestinal lymphatic transport of cyclosporine A has been reported but appears limited and formulation-dependent.^24,25^ Accordingly, the oral exposure of *ent*-verticilide despite low Caco-2 permeability may reflect favorable conformational properties, enhanced solubilization by intestinal bile salts or formulation components, underestimation of permeability due to incomplete assay recovery, or absorption pathways not fully captured by the Caco-2 model, although these mechanisms require further investigation.

The rapid NADPH-dependent disappearance of *ent*-verticilide in rat and human liver microsomes is consistent with CYP-mediated metabolism, although the individual CYP isoforms responsible were not determined. Application of the well-stirred model without correction for plasma protein or microsomal binding predicted high hepatic extraction in both species. In rats, however, the predicted hepatic clearance of approximately 63 mL/min/kg was substantially greater than the systemic clearance of 25.5 mL/min/kg measured following intravenous administration. This discrepancy may reflect extensive plasma protein binding, binding to microsomal components, and limitations associated with scaling intrinsic clearance without incorporating binding corrections. The microsomal findings should therefore be interpreted as strong evidence of metabolic liability rather than as precise quantitative predictions of in vivo hepatic clearance.

Following intravenous administration in rats, *ent*-verticilide exhibited extensive tissue distribution, moderate systemic clearance, and a terminal half-life of approximately 5 h. Systemic clearance was approximately 36% of assumed rat hepatic blood flow, indicating that clearance was not blood-flow limited despite rapid microsomal turnover. This postabsorptive profile is consistent with our previous proof-of-concept study in a murine model of catecholaminergic polymorphic ventricular tachycardia, in which intraperitoneal administration achieved sufficient systemic exposure for target engagement,^14^ suggesting that *ent*-verticilide can maintain meaningful exposure once it reaches circulation. Notably, its absolute oral bioavailability of approximately 16% represents substantial exposure for a peptide-derived macrocycle of this size. Conventional therapeutic peptides generally exhibit very low oral bioavailability without specialized structural or formulation strategies, including oral semaglutide (∼0.8%),^26^ whereas cyclic peptides such as cyclosporine (∼20% or greater) and recently optimized cyclic peptides (∼18–27% in rats) demonstrate that conformationally constrained macrocycles can achieve appreciable oral exposure.^27,28^ Thus, the oral bioavailability of *ent*-verticilide supports further development of this scaffold while leaving opportunities for structural, metabolic, and formulation optimization.

Rapid NADPH-dependent microsomal metabolism, together with the marked effects of ABT pretreatment, identifies CYP-mediated metabolism as an important limitation to *ent*-verticilide exposure. ABT substantially increased portal and systemic *C_mox_* and *AUC_O—∞_* while reducing apparent oral clearance (*Ci/F*), consistent with increased presystemic availability through reduced intestinal and/or hepatic first-pass metabolism as well as decreased postabsorptive metabolic clearance. A similar effect has been reported for cyclosporine, for which CYP inhibition can increase oral bioavailability while reducing systemic clearance.^29^ Exploratory estimates were consistent with this interpretation: *F_h_* values of approximately 0.62–0.64 suggested moderate hepatic extraction, whereas *F_a_F_g_* values of approximately 0.15–0.25 indicated a larger combined contribution from incomplete absorption and intestinal first-pass loss. However, *F_a_F_g_* cannot distinguish unabsorbed drug from intestinal metabolism, efflux, or other presystemic losses, and the ABT data therefore do not establish complete absorption or identify intestinal CYP metabolism as the predominant barrier. Likewise, the nearly superimposable portal and systemic profiles following ABT may partly reflect prolonged systemic exposure and recirculation, reducing the measurable portal–systemic concentration gradient. Collectively, these findings indicate that CYP-mediated metabolism substantially limits *ent*-verticilide exposure, while improvements in absorption may provide additional gains in oral bioavailability.

These findings have several implications for the continued preclinical development of *ent*-verticilide as an antiarrhythmic therapeutic candidate. Identification of its major metabolites, metabolic soft spots, and contributing CYP isoforms could guide structural modifications designed to reduce metabolic turnover while preserving RyR2 potency and selectivity. Solubility and permeability should be optimized in parallel with metabolic stability because incomplete absorption may remain an additional constraint on oral exposure. Lipid-based, amorphous, or other enabling formulations, together with carefully selected absorption-enhancing approaches, may improve oral bioavailability. Subcutaneous administration could provide an alternative strategy by bypassing gastrointestinal absorption and intestinal and hepatic first-pass extraction, although it would not eliminate systemic metabolic clearance. Finally, the ABT findings should be viewed as mechanistic evidence that reducing CYP-mediated metabolism can improve exposure rather than as support for coadministration of a nonspecific CYP inhibitor. Such inhibition would create substantial potential for drug–drug interactions, exposure variability, and safety concerns, particularly in patients with cardiovascular disease likely to receive multiple concomitant medications.

## CONCLUSION

The DMPK profile of the antiarrhythmic molecule *ent*-verticilide is characterized by gastrointestinal stability, extensive protein and tissue association, low apparent intestinal permeability, and rapid CYP-dependent metabolism. Following IV administration, *ent*-verticilide exhibited a large volume of distribution; however, its oral bioavailability was limited to approximately 16%. Portal and systemic pharmacokinetics measurements indicated that the fraction of the oral dose reaching the portal circulation intact (intestinal availability) was considerably lower than the fraction escaping hepatic first-pass elimination, suggesting that the major barrier to oral exposure occurs before hepatic entry. Although limited intestinal permeability likely contributes, the marked increase in portal and systemic exposure following ABT pretreatment demonstrates an important contribution of CYP-mediated presystemic metabolism. Together, these findings identify intestinal absorption and metabolism as key determinants of *ent*-verticilide exposure and suggest that strategies aimed at improving intestinal availability or reducing presystemic metabolism may substantially enhance its oral pharmacokinetic profile.

## Declaration of Generative AI Use

During the preparation of this work, the author(s) used ChatGPT-5.6 for grammatical proofreading and spelling corrections of this manuscript. The author(s) reviewed and edited the output as needed and take full responsibility for the content of the published article.

## Financial Support

This research was supported in part by the National Institutes of Health National Heart, Lung, and Blood Institute Grants [R35 HL144980] (B.C.K.), [R01 HL151223] (J.N.J., B.C.K., and W.S.A.), [R01 HL181098] (J.N.J., B.C.K., and W.S.A.), [F31 HL164064] (M.P.T.), and [F31 HL151125] (A.N.S.), the American Heart Association Arrhythmia and Sudden Death Strategically Focused Research Network [Grant 19SFRN34830019] (B.C.K), and an American Heart Association Predoctoral Fellowship grant [24PRE1243081] (T.Q.D.)

## Conflict of Interest

The authors declare no conflicts of interest.

## Data Availability

The authors declare that all the data supporting the findings of this study are available within the paper.

## CRediT Authorship Contribution Statement

**Tri. Q Do**: Conceptualization, Data curation, Formal analysis, Funding acquisition, Investigation, Methodology, Project administration, Validation, Visualization, Writing — original draft, Writing — review and editing.

**Rachel D. Crouch**: Conceptualization, Data curation, Formal analysis, Investigation, Methodology, Project administration, Validation, Visualization, Writing — original draft, Writing — review and editing.

**Kevin Wu**: Data curation, Formal analysis, Investigation, Methodology, Validation, Writing — original draft, Writing — review and editing.

**Madelaine P. Thorpe**: Data curation, Formal analysis, Funding acquisition, Investigation, Methodology, Resources, Validation, Visualization, Writing — review and editing.

**Abigail N. Smith**: Data curation, Formal analysis, Funding acquisition, Investigation, Methodology, Resources, Validation, Visualization, Writing — review and editing.

**Jeffrey N. Johnston**: Conceptualization, Data curation, Formal analysis, Funding acquisition, Investigation, Methodology, Project administration, Resources, Supervision, Validation, Visualization, Writing — review and editing.

**Wendell S. Akers**: Conceptualization, Data curation, Formal analysis, Funding acquisition, Investigation, Methodology, Project administration, Resources, Software, Supervision, Validation, Visualization, Writing — original draft, Writing — review and editing.

**Björn C. Knollmann**: Conceptualization, Data curation, Formal analysis, Funding acquisition, Investigation, Methodology, Project administration, Resources, Supervision, Validation, Visualization, Writing — review and editing.

## List of Non-Standard Abbreviations

AAG: alpha1-acid glycoprotein
ABT: 1-aminobenzotriazole
AUC: Area under the concentration versus time curve
AUMC: Area under the (time x concentration) versus time curve
Ca: calcium
Caco-2: polarized human colorectal adenocarcinoma-derived cells
CYP: Cytochrome P450
DMSO: dimethyl sulfoxide
DMPK: Drug Metabolism and Pharmacokinetics
FaSSGF: Fasted state simulated gastric fluid
FaSSIF: Fasted state simulated intestinal fluid
HBSS: Hank’s balanced salt solution
HPLC: High-performance liquid chromatography
HPV: Hepatic portal vein
HSA: Human serum albumin
JVC: Jugular vein cannulation
LC-MS/MS: Liquid chromatography tandem mass spectrometry
MgCl_2_: Magnesium Chloride
MRT: Mean residence time
NADPH: nicotinamide adenine dinucleotide phosphate tetrasodium salt
PBS: Phosphate buffered saline
QC: Quality control
RyR2: Ryanodine receptor 2
TEER: Transepithelial electrical resistance

## Notes

### Competing Interest Statement

The authors have declared no competing interest.

## Reference

1. Bers DM. Cardiac excitation-contraction coupling. Nature. 2002;415(6868):198–205. 10.1038/415198a

2. Do TQ, Knollmann BC. Inhibitors of Intracellular RyR2 Calcium Release Channels as Therapeutic Agents in Arrhythmogenic Heart Diseases. Annu Rev Pharmacol Toxicol. 2025;65(1):443–463. 10.1146/annurev-pharmtox-061724-080739

3. Batiste SM, Blackwell DJ, Kim K, et al. Unnatural verticilide enantiomer inhibits type 2 ryanodine receptor-mediated calcium leak and is antiarrhythmic. Proc Natl Acad Sci U S A. 2019;116(11):4810–4815. 10.1073/pnas.1816685116

4. Shiomi K, Matsui R, Kakei A, et al. Verticilide, a new ryanodine-binding inhibitor, produced by Verticillium sp. FKI-1033. J Antibiot (Tokyo). 2010;63(2):77–82. 10.1038/ja.2009.126

5. Gochman A, Do TQ, Kim K, et al. ent-Verticilide B1 inhibits type 2 ryanodine receptor channels and is antiarrhythmic in Casq2−/− mice. Mol Pharmacol. 2024;10.1124/molpharm.123.000752

6. Nielsen DS, Shepherd NE, Xu W, Lucke AJ, Stoermer MJ, Fairlie DP. Orally Absorbed Cyclic Peptides. Chem Rev. 2017;117(12):8094–8128. 10.1021/acs.chemrev.6b00838

7. Fouche M, Schafer M, Berghausen J, et al. Design and Development of a Cyclic Decapeptide Scaffold with Suitable Properties for Bioavailability and Oral Exposure. ChemMedChem. 2016;11(10):1048–59. 10.1002/cmdc.201600082

8. Zientek M, Jiang Y, Youdim K, Obach RS. In vitro-in vivo correlation for intrinsic clearance for drugs metabolized by human aldehyde oxidase. Drug Metab Dispos. 2010;38(8):1322–7. 10.1124/dmd.110.033555

9. Carlile DJ, Zomorodi K, Houston JB. Scaling factors to relate drug metabolic clearance in hepatic microsomes, isolated hepatocytes, and the intact liver: studies with induced livers involving diazepam. Drug Metab Dispos. 1997;25(8):903–11.

10. Barter ZE, Bayliss MK, Beaune PH, et al. Scaling factors for the extrapolation of in vivo metabolic drug clearance from in vitro data: reaching a consensus on values of human microsomal protein and hepatocellularity per gram of liver. Curr Drug Metab. 2007;8(1):33–45. 10.2174/138920007779315053

11. Davies B, Morris T. Physiological parameters in laboratory animals and humans. Pharm Res. 1993;10(7):1093–5. 10.1023/a:1018943613122

12. Obach RS. Prediction of human clearance of twenty-nine drugs from hepatic microsomal intrinsic clearance data: An examination of in vitro half-life approach and nonspecific binding to microsomes. Drug Metab Dispos. 1999;27(11):1350–9.

13. Di L, Feng B, Goosen TC, et al. A perspective on the prediction of drug pharmacokinetics and disposition in drug research and development. Drug Metab Dispos. 2013;41(12):1975–93. 10.1124/dmd.113.054031

14. Blackwell DJ, Smith AN, Do T, et al. In Vivo Pharmacokinetic and Pharmacodynamic Properties of the Antiarrhythmic Molecule ent-Verticilide. J Pharmacol Exp Ther. 2023;385(3):205–213. 10.1124/jpet.122.001455

15. de Montellano PRO. 1-Aminobenzotriazole: A Mechanism-Based Cytochrome P450 Inhibitor and Probe of Cytochrome P450 Biology. Med Chem (Los Angeles). 2018;8(3)10.4172/2161-0444.1000495

16. Matsuda Y, Konno Y, Satsukawa M, et al. Assessment of intestinal availability of various drugs in the oral absorption process using portal vein-cannulated rats. Drug Metab Dispos. 2012;40(12):2231–8. 10.1124/dmd.112.048223

17. Matsuda Y, Konno Y, Hashimoto T, et al. Quantitative assessment of intestinal first-pass metabolism of oral drugs using portal-vein cannulated rats. Pharm Res. 2015;32(2):604–16. 10.1007/s11095-014-1489-x

18. Laznicek M, Laznickova A. The effect of lipophilicity on the protein binding and blood cell uptake of some acidic drugs. J Pharm Biomed Anal. 1995;13(7):823–8. 10.1016/0731-7085(95)01504-e

19. Brandon AM, Baginski SR, Peet C, et al. Log D(7.4) and plasma protein binding of synthetic cannabinoid receptor agonists and a comparison of experimental and predicted lipophilicity. Drug Test Anal. 2024;16(9):1012–1025. 10.1002/dta.3621

20. Thorpe MP, Smith AN, Blackwell DJ, et al. The backbone constitution drives passive permeability independent of side chains in depsipeptide and peptide macrocycles inspired by ent-verticilide. Chem Sci. 2024;15(36):14977–87. 10.1039/d4sc02758b

21. Thorpe MP, Hopkins CR, Johnston JN. End-to-End Backbone Cyclization Enhances Passive Permeability of bRo5 Oligomeric Depsipeptides with Nonlinear Size Dependence. ACS Med Chem Lett. 2025;16(4):638–645. 10.1021/acsmedchemlett.5c00037

22. Smith AN, Blackwell DJ, Knollmann BC, Johnston JN. Ring Size as an Independent Variable in Cyclooligomeric Depsipeptide Antiarrhythmic Activity. ACS Med Chem Lett. 2021;12(12):1942–1947. 10.1021/acsmedchemlett.1c00508

23. Do TQ, Blackwell DJ, Smith AN, et al. Introduction of a single carboxylic acid converts the cyclic oligomeric depsipeptide ent-verticilide from a ryanodine receptor 2 (RyR2) inhibitor to RyR2 activator. Mol Pharmacol. 2025;107(12):100086. 10.1016/j.molpha.2025.100086

24. David W, Foley IB, Patrick D. Bailey, David Meredith. A cyclosporine derivative is a substrate of the oligopeptide transporter PepT1. MedChemComm. 2016;7(5):999–1002. 10.1039/C5MD00575B

25. Ueda CT, Lemaire M, Gsell G, Nussbaumer K. Intestinal lymphatic absorption of cyclosporin A following oral administration in an olive oil solution in rats. Biopharm Drug Dispos. 1983;4(2):113–24. 10.1002/bdd.2510040203

26. Overgaard RV, Navarria A, Ingwersen SH, Baekdal TA, Kildemoes RJ. Clinical Pharmacokinetics of Oral Semaglutide: Analyses of Data from Clinical Pharmacology Trials. Clin Pharmacokinet. 2021;60(10):1335–1348. 10.1007/s40262-021-01025-x

27. Hill TA, Lohman RJ, Hoang HN, et al. Cyclic Penta- and Hexaleucine Peptides without N-Methylation Are Orally Absorbed. ACS Med Chem Lett. 2014;5(10):1148–51. 10.1021/ml5002823

28. Merz ML, Habeshian S, Li B, et al. De novo development of small cyclic peptides that are orally bioavailable. Nat Chem Biol. 2024;20(5):624–633. 10.1038/s41589-023-01496-y

29. Gomez DY, Wacher VJ, Tomlanovich SJ, Hebert MF, Benet LZ. The effects of ketoconazole on the intestinal metabolism and bioavailability of cyclosporine. Clin Pharmacol Ther. 1995;58(1):15–9. 10.1016/0009-9236(95)90067-5

